# MetagenBERT: a Transformer Architecture using Foundational DNA Read Embedding Models to enhance Disease Classification

**DOI:** 10.1101/2025.05.06.652444

**Authors:** Gaspar Roy, Eugeni Belda, Edi Prifti, Yann Chevaleyre, Jean-Daniel Zucker

## Abstract

Microbial ecosystems constitute complex yet information-rich environments whose characterization is crucial for understanding host health and disease. Among them, the human gut microbiome has emerged as a key "super-integrator", owing to its dense interactions with host physiology and its established associations with a wide spectrum of pathologies. Driven by advances in high-throughput sequencing technologies and the continuous decline in associated costs, metagenomic studies have expanded exponentially, generating massive amounts of sequencing data and opening new avenues for data-driven disease modeling. Conventional approaches to microbiome analysis predominantly rely on the alignment of DNA sequencing reads against reference databases to infer microbial composition and profiling at the species level. While effective, these methods are inherently constrained by reference bias and limited taxonomic resolution. Recent advances in artificial intelligence—particularly in Natural Language Processing (NLP) offer new methodological perspectives for metagenomic data representation. In this study, we present MetagenBERT, a Transformer-based framework to embed metagenomes that relies on the foundational models DNABERT-2 and DNABERT-S for the embedding of DNA sequencing reads. Our approach encodes gut microbiome metagenome in a taxonomy-agnostic manner, enabling direct downstream application to disease classification tasks. We demonstrate that MetagenBERT reaches similar performance to state-of-the-art abundance-based models for cirrhosis prediction and surpasses them in the more challenging context of type 2 diabetes. Furthermore, we introduce an alternative representation of metagenomes based on read-level embeddings aggregated into abundance vectors, demonstrating their complementarity with conventional species-level abundance metrics.

## 1. Introduction

The human gut microbiome, consisting of bacteria, fungi, viruses, archaea, and eukaryotes, outnumbers human cells tenfold Ferranti et al., 2014. Its diversity and composition are critical indicators of patient health status Pflughoeft and Versalovic, 2012 Le Chatelier et al., 2013 Qin et al., 2014. Next-Generation Sequencing (NGS) has enabled cost-effective analyses of microbial ecosystems, advancing fields like metagenomics, which profiles the whole DNA from a sample by generating typically millions of short sequences per sample (100 to 300 bases) MetaHIT Consortium et al., 2010. While long-read sequencing is emerging Oehler et al., 2023, allowing to retrieve reads of thousands if not millions of bases, short-read methods remain dominant due to cost-effectiveness and their ability to estimate quantification profiles.

Traditional bioinformatics methods for microbiome sample analysis involves identifying the species it contains, computing their relative abundance tables, and eventually linking the species to health conditions. They mostly rely on large reference catalogs, making them computationally expensive AltschuP et al., n.d. A typical pipeline is described in Figure 1. To reduce dependence on reference databases, Deep Learning (DL) techniques—including sequence classification via TetraNucleotide Frequency (TNF) Sharma et al., 2020Nissen et al., 2021 or Natural Language Processing (NLP)Mock et al., 2021Liang et al., 2020Roy et al., 2024 can be used, with the objective to automatically identify abstract representations.

**Figure 1.**
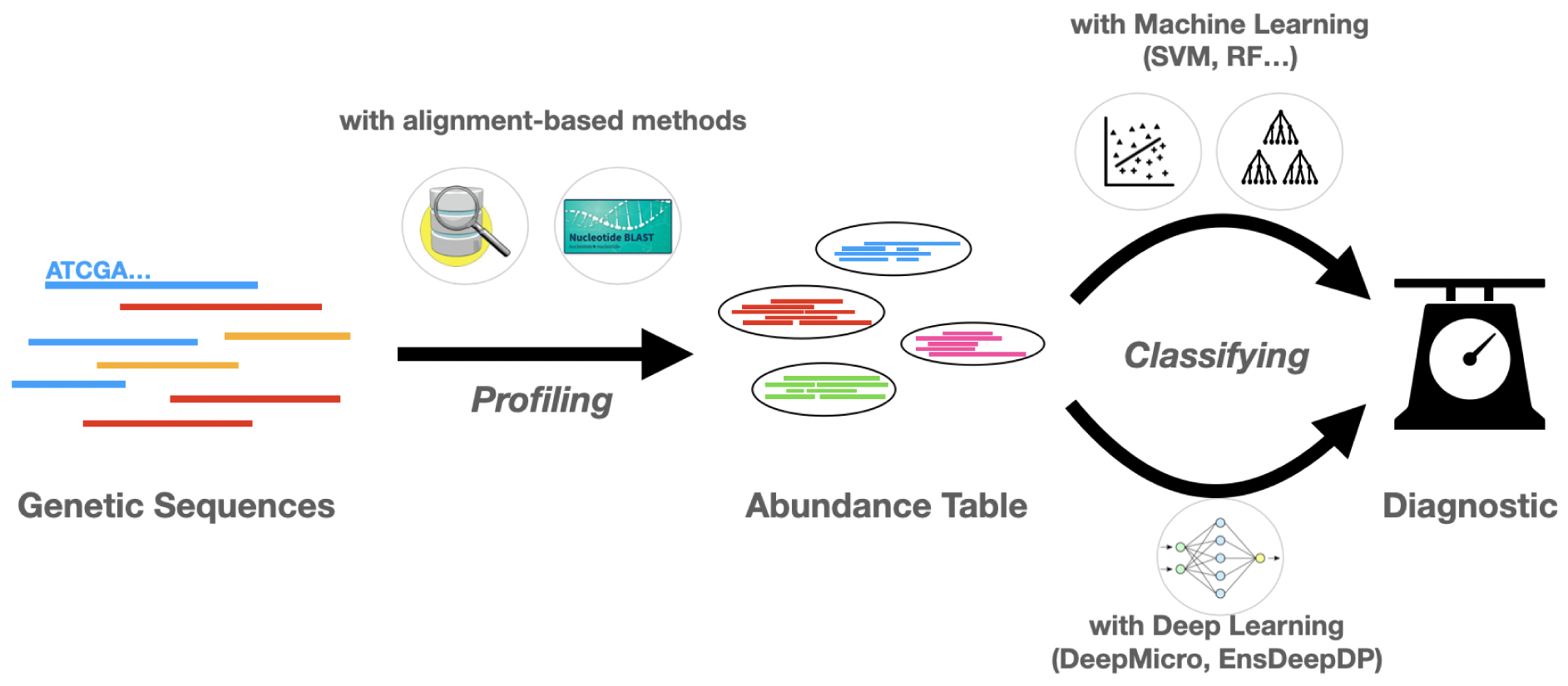
Typical bioinformatics pipeline for metagenomics analyses. The widely used approach to perform disease prediction from WGS data is to bin each read into a reference catalog of genes or species. This results in a data table of presence or relative abundance, which can be further used to classify samples in disease groups. This table can then be analyzed to perform classification.

Recently, Large Language Models (LLMs) have been increasingly applied in genomics. Foundation models such as Nucleotide Transformer Dalla-Torre et al., 2025 and DNABERT-2 Zhou et al., 2024a have demonstrated promising performance in various classical gene analysis tasks. More specifically, DNABERT-2 has been further fine-tuned into DNABERT-S Zhou et al., 2024b using contrastive learning to generate embeddings—mathematical vector representations of sequences—by pulling together sequences from closely related species while pushing apart those from more distant species. This approach enhances sequence-to-species grouping (also known as binning).

Meanwhile, MetaTransformer Wichmann et al., 2023, an attention-based model without pre-training, has exhibited interesting performance in sequence classification tasks, highlighting the potential of transformer-based models in genomics.

In order to link microbiome composition to patient health status, different DL approaches such as PopPhy-CNN Reiman et al., 2020, MML4Microbiome Lee and Rho, 2022, EnsDeepDP Shen et al., 2022 and DeepMicro Oh and Zhang, 2020 have been developed to extract abundance features, but still struggle with the high-dimensional, sparse nature of abundance tables, risking overfitting. Solutions like data augmentation Mulenga et al., 2021 or simulation can improve training but may reduce diversity Sczyrba et al., 2017. Some methods bypass abundance tables altogether, using direct sequence embeddings and averaging those by species, as in Metagenome2Vec Queyrel et al., 2020.

However, studies have identified several limitations in using species composition for predicting metagenomic disease associations. First, species detection relies heavily on reference catalogs, which, despite their continuous expansion, still fails to capture a substantial portion of microbiome diversity Thomas and Segata, 2019. Additionally, the choice of taxonomic resolution significantly impacts prediction accuracy. Research has demonstrated that individuals from the same species can exhibit varying, and sometimes even opposing, effects on disease development. Consequently, alternative approaches, such as functional guild-based representations, have been proposed to improve the representation of metagenomic samples Wu et al., 2021.

A key challenge in metagenomic analysis is the nature of metagenomic data, which consists of vast amounts of short reads—often numbering in the tens of millions per sample—while datasets typically comprise only a few hundred samples. As a result, metagenomic datasets are highly complex, requiring feature extraction from a limited number of examples. This imbalance makes it difficult to identify meaningful patterns that generalize well for classification tasks while mitigating the risk of overfitting. Calle, 2019

For these reasons, we proposed to explore the feasibility an end-to-end and species-agnostic approach, based on Transformers architectures, leveraging powerful embeddings and aggregation techniques for improved microbiome classification and disease prediction. These aggregations also propose a structure comparable to abundance tables but relying on a species-agnostic approach that captures information different from the classic abundance table.

## 2. Materials and Methods

The primary objective of our approach is to develop a method that is both end-to-end and independent of reference catalogs. This method directly processes all DNA reads from a metagenome (after preprocessing) and performs classification without reducing the information to species-level proportions. The underlying idea is that DNA reads contain valuable information beyond species identity, and this information can be effectively extracted using DNA-based large language models.

### 2.1. Datasets

To train and evaluate our method, we used two metagenomic datasets, also employed in the MetaML study Pasolli et al., 2016. These datasets are related to two clinical conditions, Liver Cirrhosis cirr and Type 2 Diabetes t2d. Both have a comparable proportion of disease and control samples. More information on these datasets can be found in Table 1. We downloaded the raw fastq files from EBI (ERP005860) for cirrhosis and NCBI ( SRA045646 and SRA050230) for diabetes and cleaned them using fastp with default parameters as illustrated in Chen, 2023.

**Table 1.** Summary of the two datasets used along with computational information.

|  | Number of Samples (H/D) | Mean Number of reads per sample | Standard Deviation | Storage | Embedding Time (hours) | Storage of Embeddings | Global Clustering Time |
| --- | --- | --- | --- | --- | --- | --- | --- |
| Cirrhosis | 232 (114/118) | 36,823M | 23,396M | 1.1T | 12h | 25,201 T | 18h |
| Type 2 Diabetes | 344 (174/170) | 34,467M | 12,214M | 808G | 26h | 34,977 T | 30h |

### 2.2. Pipeline for Metagenome Embedding

Our achitecture consists of several steps detailed in the next subsections. First, the reads from a sample are embedded using a Large Language Model (LLM). Then, depending on the method developed later, this information will either be pooled to form a single vector, the metagenome embedding, or some specific vectors will be selected to represent the metagenome as a smaller bag of vectors, thus drastically reducing its complexity. Keeping in mind that a metagenome is composed of millions of reads, it is challenging to compress the information from millions of vectors to a single one, both for computational reasons and because of the risk of information loss during aggregation.

#### 2.2.1. Read embedding through Large Language Models

The first step of our pipeline consists in transforming the single reads into embeddings. This can be approached by using LLM models trained on DNA data. We used DNABERT-2 Zhou et al., 2024a, a model based on the MosaicBERT Portes et al., n.d. architecture, which learns the DNA features by performing Masked Language Modeling on genomic data extracted from various species. For each model, and each sample of our datasets, we embedded the totality of the available sequences, running inference with a batch size of 40000. The embeddings dimension was 768, and the maximum length of a sequence in token was 60 tokens. We then applied average pooling of the embeddings on the token axis, therefore representing the sequences as vectors of dimension 768. Finally, we also used DNABERT-S Zhou et al., 2024b, a fine-tuned version of DNABERT-2 that was explicitly trained to differentiate sequences from different species through contrastive learning, thus learning to generate close embeddings for reads from the same species and more different embeddings for reads originating from different species. Our motivation in focusing in these two architectures, was to obtain both species-specific embeddings with DNABERT-S and more general embeddings with DNABERT-2. All the following steps were applied to both DNABERT-2 and DNABERT-S embedded metagenomes separately. We employed nodes of the Jean-Zay cluster, composed of 96 A100 GPUs to infer the embeddedings of the sequences from both datasets, with a batch size of 40000 reads.

#### 2.2.2. Aggregating Read Information

We can consider the read embeddings as local features of our metagenome, we have then used aggregating, sub-sampling and clustering methods in order to represent the global structure of the microbiome.

#### 2.2.3. Simple Aggregation Method

As a baseline, we used a simple aggregation method that computes the mean of all embeddings. However, this approach is inherently oversimplifying and leads to significant information loss. For instance, reads with opposing embeddings in certain dimensions may cancel each other out, erasing meaningful variations in the data. Additionally, this method is likely to be biased toward highly represented species, as their read embeddings may cluster more closely together, further limiting the effectiveness of the representation.

**Figure 2.**
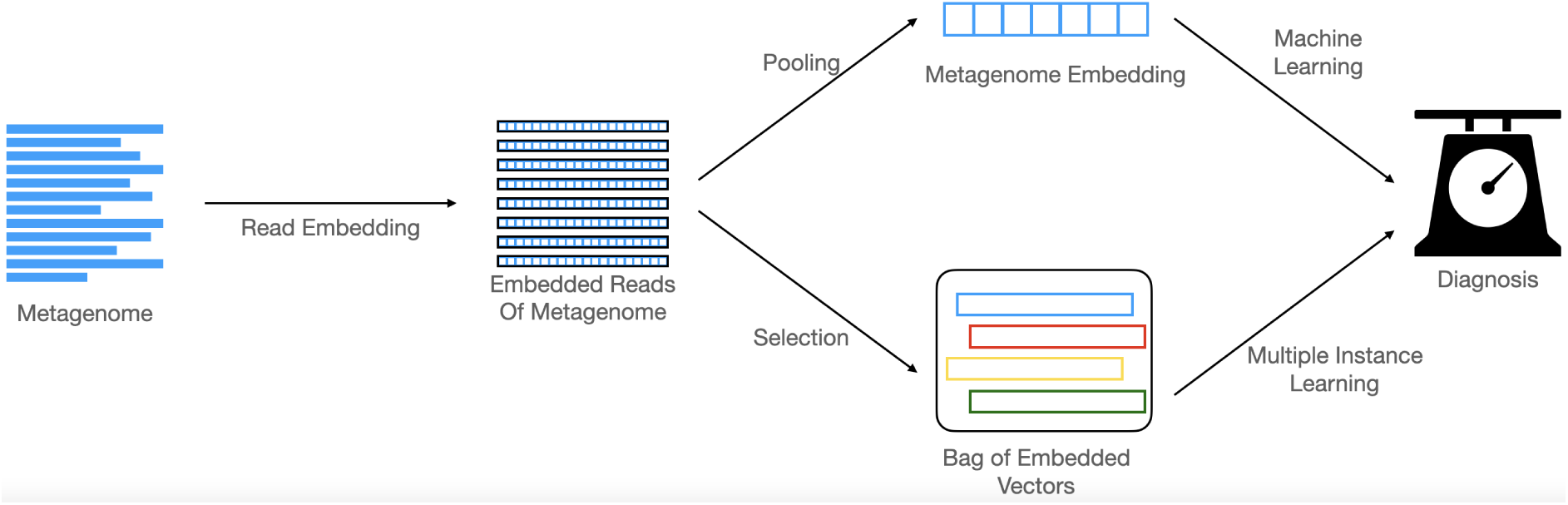
The Architecture of our pipeline : metagenomic samples are processed one at a time and their reads embedded by a Transformer model. The metagenome is then represented as a set of embeddings of reads. In order to use it for classification, its dimension is then either reduced to a single vector through aggregation (3), or to a smaller set of vectors through clustering (4) or sampling operations (5). The resulting object can then be used for classification respectively with Machine Learning algorithms or DeepSets Zaheer et al., 2018

**Figure 3.**
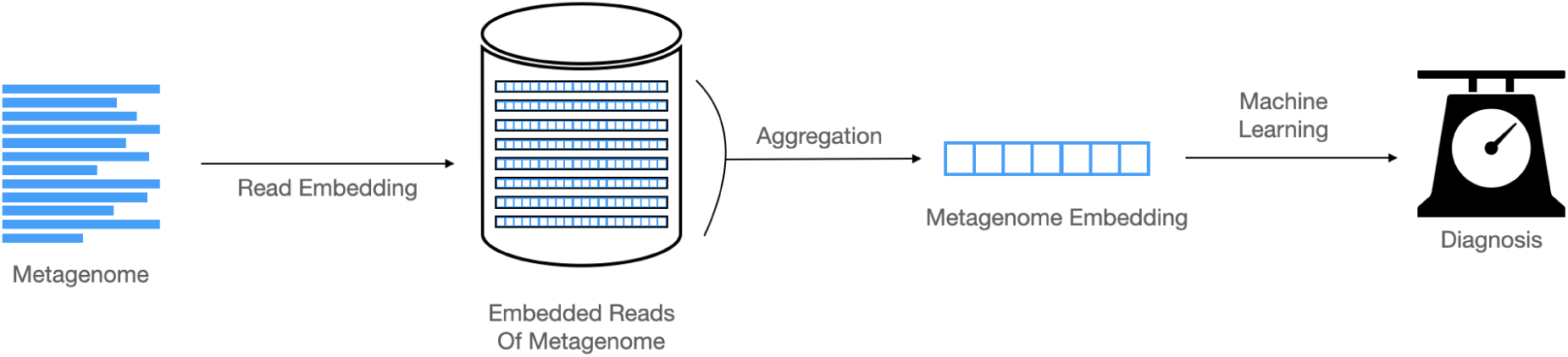
The Simple Aggregation Architecture : the set of embedded reads is transformed into one vector by a simple mean operation, then used for classification.

#### 2.2.4. Clustering Methods

In order to reduce the size of our metagenome representation while keeping different types of information, we decided to regroup our read embeddings in clusters, each cluster representing a part of the metagenome. For clustering, we used FAISS Kmeans implementation for rapid, efficient, GPU-optimized clustering Douze et al., 2024Johnson et al., 2019. We repeated the experiment with a number of clusters ranging from 16 to 16384 (increasing by a factor of 2 each time). The clustering returns two elements : the centroids of each cluster and the assignment of each vector populating the clusters. These assignements can then be used to calculate the proportion of reads in each cluster for each metagenome. We have used this method to develop two different types of clustering pipelines.

Local Clustering. When using the method we called "Local Clustering", or MetagenBERT-Local, we performed the KMeans algorithm independantly on each sample of each dataset. A subsample of the reads is used to train the KMeans, then each vector is assigned to its corresponding cluster. The centroids of each cluster are then retrieved to represent the metagenome. In this case, the metagenome is represented by an unordered set of vectors of dimensions number of clusters*embedding size. In this case, each clustering being independent on the sample, calculating the abundance is not useful, for the i-th cluster in the first metagenome is not linked at all to the i-th cluster of the second metagenome. Only the centroids of the clusters are relevant here to represent a metagenome.

Global Clustering. The method designated as "Global Clustering", or MetagenBERT-Glob, although also relying on clustering, is different both in its goal and final representation. The idea here is to create a new abundance table representing the microbiome, but based on our embeddings instead of being based on species. To achieve this, we need the different clusters to represent the same parts of the microbiome across all samples from a dataset. To do so, we train our clustering method with reads from 90% of the samples in the dataset (and leave the 10 last percent as holdout). We use 240,000 reads from each sample in the cirrhosis dataset and 180,000 from each sample in the Type 2 Diabetes dataset (for memory considerations), so approximately 1% per sample. This ensures a good enough coverage from every sample in the dataset. Once the KMeans is trained, we take each sample one by one and assign all its reads to their corresponding cluster using nearest neighbors. In this situation, as opposed to local clustering method, samples are not represented by their centroids (as those are common to the whole dataset), but rather by a unique vector : the abundance of each cluster.

Combining Global Clustering Abundance with Species Abundance. In order to compare our abundance vector to species-based abundance, we retrieved the abundance vectors of each sample in the dataset and trained a model for disease prediction with the following configurations : by using species-abundance alone, by using our clustering abundance alone for each number of clusters and by using a concatenation of both for each number of clusters. This experiment allows to compare our method to the baseline, but also to see if the combination of both information makes the results better. If so, this might mean that both methods extract different features from the metagenome.

#### 2.2.5. Subsampling Method

The subsampling method is a simple baseline method that we tested in order to make sure our local clustering method was relevant. To prove that our clusters were significant, and their centroids were good representatives of the different parts of our metagenome, we also tried selecting random reads from each of our samples, choosing as many as we had clusters and treating these reads as if they were the centroids of our clusters. A sample is then represented by a set of vectors and its dimensions are number of samples * embedding size. We expect to obtain worse results with this method than with the local clustering method, thus showing that clustering efficiently captures different relevant parts of our samples.

### 2.3. Classification from Metagenome Embedding

Once the aggregation methods have been applied, the resulting embedding can be utilized to train a classification model. Two distinct scenarios can be considered based on the representation of the metagenome.

- Single-Vector Representation: When employing Simple Aggregation or Global Clustering, the metagenome is represented as a single feature vector. This allows for the application of standard machine learning algorithms, such as Lasso regression, Random Forests, or Multi-Layer Perceptrons (MLP) mlp.
- Set-Based Representation (Multiple Instance Learning): On the other hand, the remaining aggregation methods, local clustering and subsampling, yield a representation in which the metagenome is characterized as an unordered set of feature vectors. This formulation aligns with the Multiple Instance Learning (MIL) framework Babenko, n.d., wherein a sample is represented by a collection of instances rather than a single feature vector.

**Figure 4.**
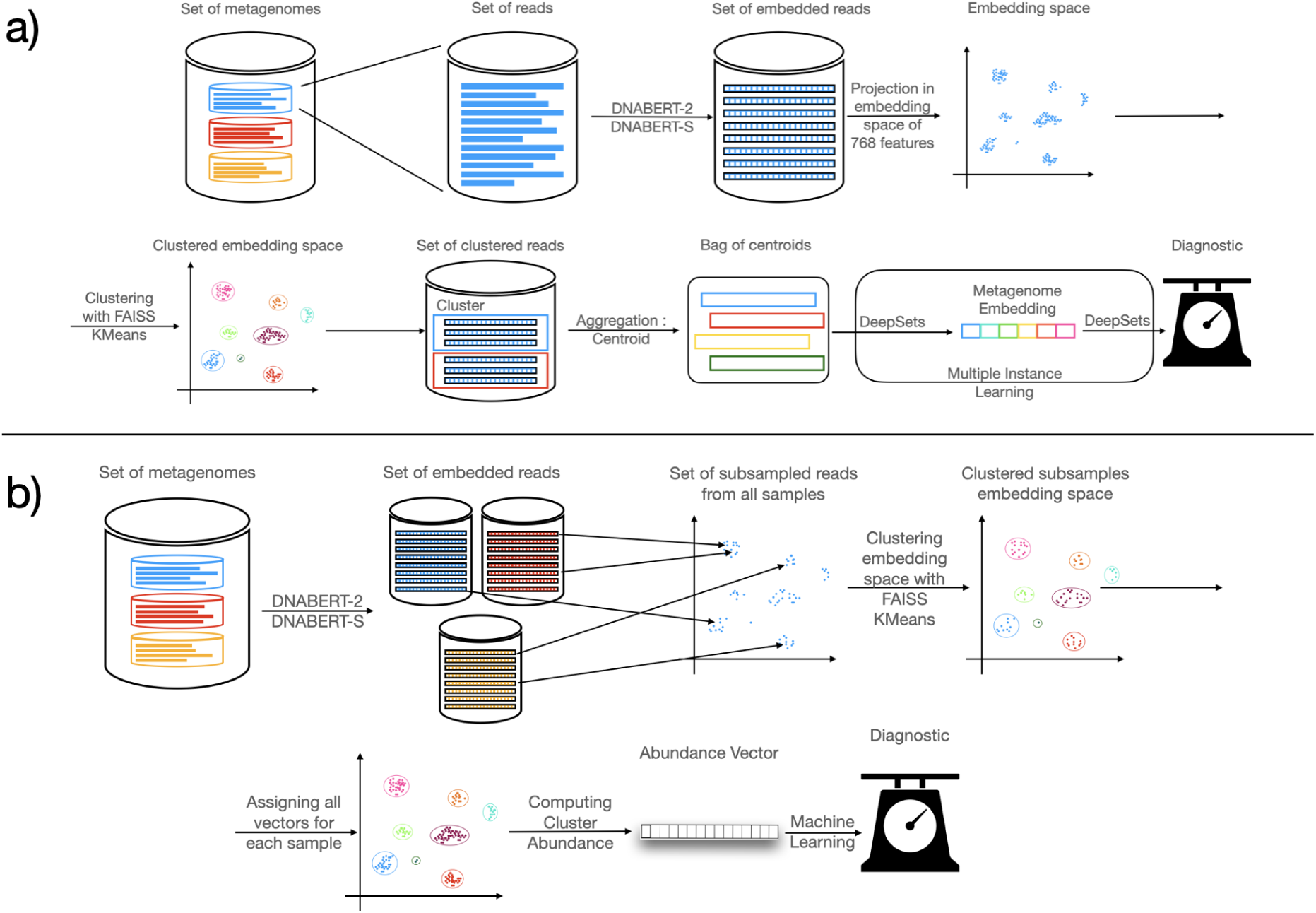
The Clustering Architectures : a) represents the Local Clustering method : each sample is independently considered as a set of embedded reads and clusterized using KMeans algorithm from FAISS implementation. The centroid of each cluster is then kept as a representative to produce a new subsample bag that is classified with DeepSets. b) represents the Global Clustering method : a subsample of each embedded metagenome is used to train a KMeans common to all the chosen dataset. Every embedding of each sample is then assigned to its cluster, thus creating a new abundance vector based on embeddings rather than species used for classification.

**Figure 5.**
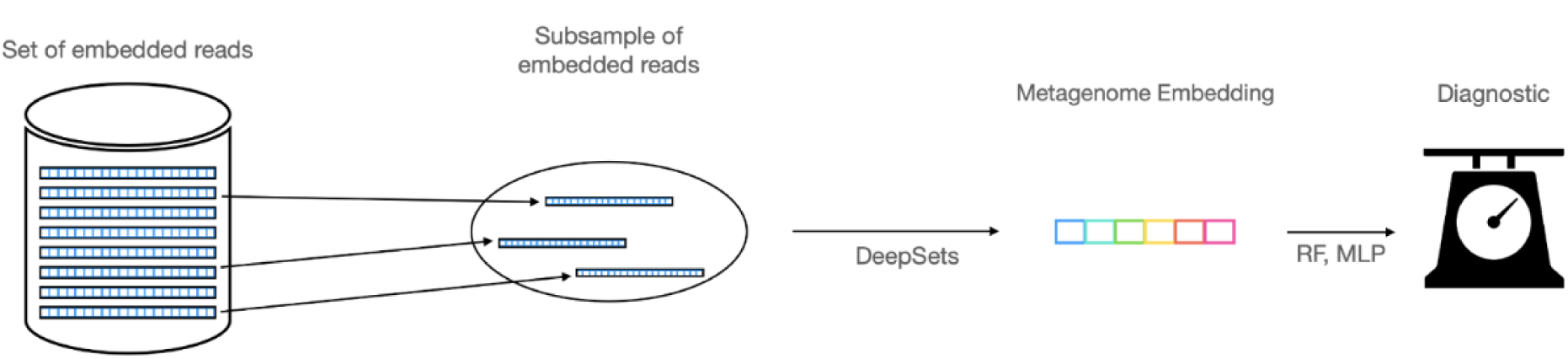
The Subsampling Method : a random subsample is drawn from the set of reads, thus obtaining a smaller set that we classify using a DeepSets network.

To address this MIL problem, we adopt the DeepSets architecture Zaheer et al., 2018. DeepSets consists of two neural networks, *ϕ* and *ρ*, separated by a permutation-invariant pooling layer. The first network, *ϕ*, processes each vector independently to extract relevant features. These extracted features are then aggregated using the pooling layer, producing a global representation of the metagenome. The second network, *ρ*, subsequently analyzes this global representation and performs the final disease classification.

In both cases, we performed 10-fold cross validation and computed the standard classification error for each experiment.

### 2.4. Global Clustering Pipeline with 10% data

A metagenomic sample consists of a vast collection of diverse reads, and our approach necessitates a continuous balance between representativity and information aggregation. Embedding and clustering all reads within each metagenomic sample demand substantial computational time and resources. To improve efficiency and reproducibility, we explored a streamlined version of our method by utilizing only a fraction of the data (10%) to assess whether comparable results could be achieved relative to the full dataset : our global clustering model is trained using less than 10% of the available data—specifically, 240,000 reads per sample for the cirrhosis dataset and 180,000 reads per sample for the diabetes dataset. Subsequently, only 10% of the reads from each sample are assigned to the resulting clusters, and cluster proportion vectors are computed based on this subset, rather than on the full set of reads.

### 2.5. Clusters Taxonomic Analysis

Our results suggest that the clusters identified by our method capture distinct dynamics compared to those inferred from species composition. To further investigate this, we analyzed the content of clusters in the T2D dataset derived from DNABERT2 under the global clustering setting with 2048 clusters. For each sample, we generated FASTA files for the 2048 clusters by mapping read assignments back to the original sample FASTA files. We then used Centrifuge v1.0.4 to taxonomically bin individual cluster reads to reference genome sequences from the UHGC1.0 catalog. Reads assigned to species-level representative genomes were retained (min. 10 reads), from which cluster shannon entropy was computed as:

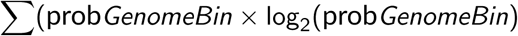

Where probGenome Bin is the ratio of the number of reads assigned to the Genome by Centrifuge divided by the total number of reads assigned to species-level representative genomes. From this, the mean value of the entropy for the 2048 clusters in each sample were retained as metric describing the complexity of the sample in terms of sequence embeddings.

## 3. Results

### 3.1. Baseline Classification using Simple Aggregation

As a baseline, we performed classification on cirrhosis and diabetes datasets using the simple mean of all vector representations. The results, presented in Table 2, indicate that while this approach provides some insights for cirrhosis, its performance is significantly lower compared to state-of-the-art methods. This outcome is expected, as the aggregation process leads to substantial information loss and fails to catch the diversity of the metagenome. Although the performances in classifying Type 2 Diabetes are weaker, they are better than could be expected when considering the difficulty of the task and the State of the Art results.

**Table 2.** Performance of MetagenBERT-Aggreg on cirrhosis and type 2 diabetes using DNABERT-2 and DNABERT-S embeddings. Metrics are Accuracy and AUC, in parenthesis is given the standard classification error.

|  |  | Cirrhosis | Type 2 Diabetes |
| --- | --- | --- | --- |
| Accuracy | DNABERT-2 | 80.63% (2.56) | 69.91% (2.31) |
|  | DNABERT-S | 82.3% (2.44) | 70.94% (2.26) |
| AUC | DNABERT-2 | 0.8797 (0.0210) | 0.7711 (0.0219) |
|  | DNABERT-S | 0.8827 (0.0206) | 0.7860 (0.0232) |

### 3.2. Clustering methods allow valuable insights on microbiome dynamics

#### 3.2.1. Local clustering methods compete with State-of-the-Art

As shown in Figure 6, the local clustering method yields results that are competitive with state-of-the-art approaches such as MetaML, PopPhyCNN, DeepMicro, MML4Microbiome, and EnsDeepDP on the cirrhosis dataset. While performance is weaker for a small number of clusters, it improves as the number of clusters increases—enhancing the representativity of the sample until it reaches a plateau. The highest performance is achieved with 2048 clusters, with an AUC of 0.929 and an accuracy of 89.24%).

**Figure 6.**
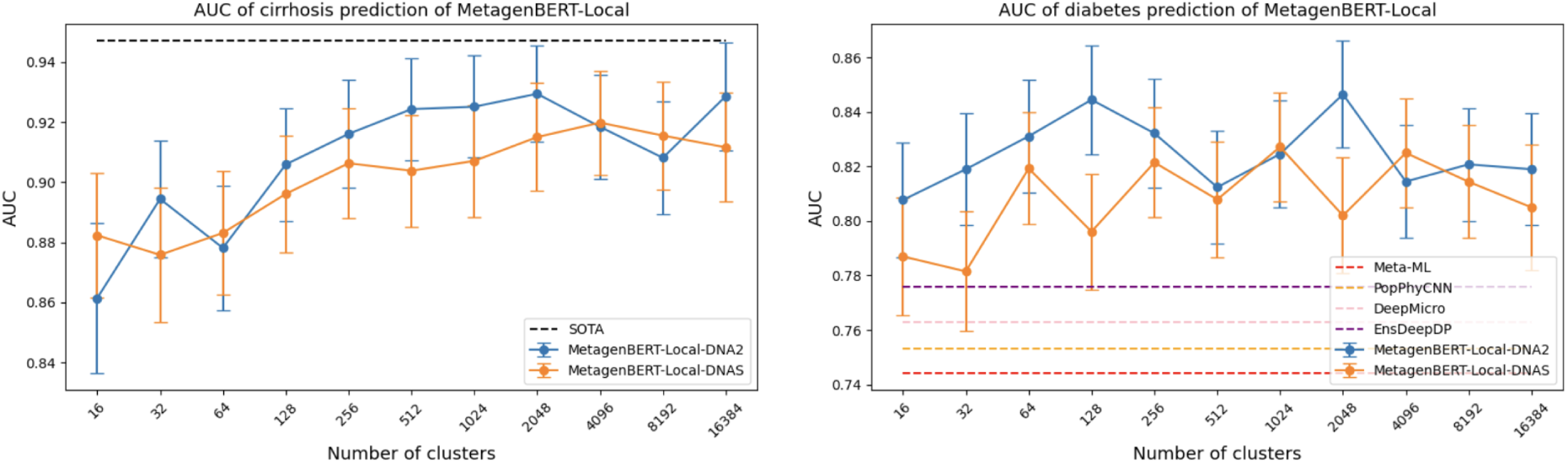
Performance Comparison between various SOTA models and our MetagenBERT local clustering classification. Local clustering results in comparable performance to SOTA models on cirrhosis and outperform them on Type 2 Diabetes. For cirrhosis, figure shows the importance of a high number of clusters in prediction quality.

For the more challenging task of diabetes prediction, the best results are achieved with a smaller number of clusters (128), generally outperforming other methods. In the best case, our approach attains 80.28% accuracy and an AUC of 0.844. These findings suggest two nonexclusive conclusions. First, the clusters generated by our method provide a different representation from species-based approaches, which may be better suited for difficult classification tasks where species-level information is insufficient. Second, the Transformer-based embeddings of reads, found in our centroids, contain relevant features for classification, supporting the hypothesis that Transformers can capture information relevant to the functional role of sequences in microbiome dynamics.

In general, we see the results obtained when using DNABERT-S are slightly weaker than with DNABERT-2, although very close. We can only assume this means that DNABERT-2 creates more general embeddings in which some information different from the ones used to represent the specie of origin are contained.

#### 3.2.2. Global Clustering methods present a different source of information on microbiome dynamics than specie abundance

As previously mentioned, we compared different configurations of our method against state-of-the-art results, including classification using species abundance alone, our global clustering abundance method alone, and a combination of both. We used a simple LASSO classifier, whereas other methods leverage additional sources of information — such as phylogeny in PopPhyCNN Reiman et al., 2020 and multimodal data in MML4Microbiome Lee and Rho, 2022 — or more complex learning strategies, like CNNs or ensemble learning in EnsDeepDP Shen et al., 2022.

Our results indicate that, for a low number of clusters, the global clustering abundance method underperforms due to an insufficient number of features to compete with species-based abundance. However, as the number of clusters increases, it ultimately outperforms species abundance and achieves performance comparable to state-of-the-art methods. These findings suggest that our global clustering approach effectively captures important microbiome dynamics.

Additionally, using abundance features alone, rather than centroids, demonstrates that our method’s performance is not solely reliant on the expressive power of Transformer-extracted centroids. Instead, the way reads are distributed within the embedding space also carries valuable information for disease prediction. Lastly, we observe that combining global clustering abundance with species abundance almost always improves classification performance, indicating that these two feature vectors encode different and potentially complementary information.

We therefore conclude that MetagenBERT-Glob may represent a novel source of insight into microbiome data and can be used alongside species abundance to enhance our understanding of the differences between healthy and diseased states.

**Figure 7.**
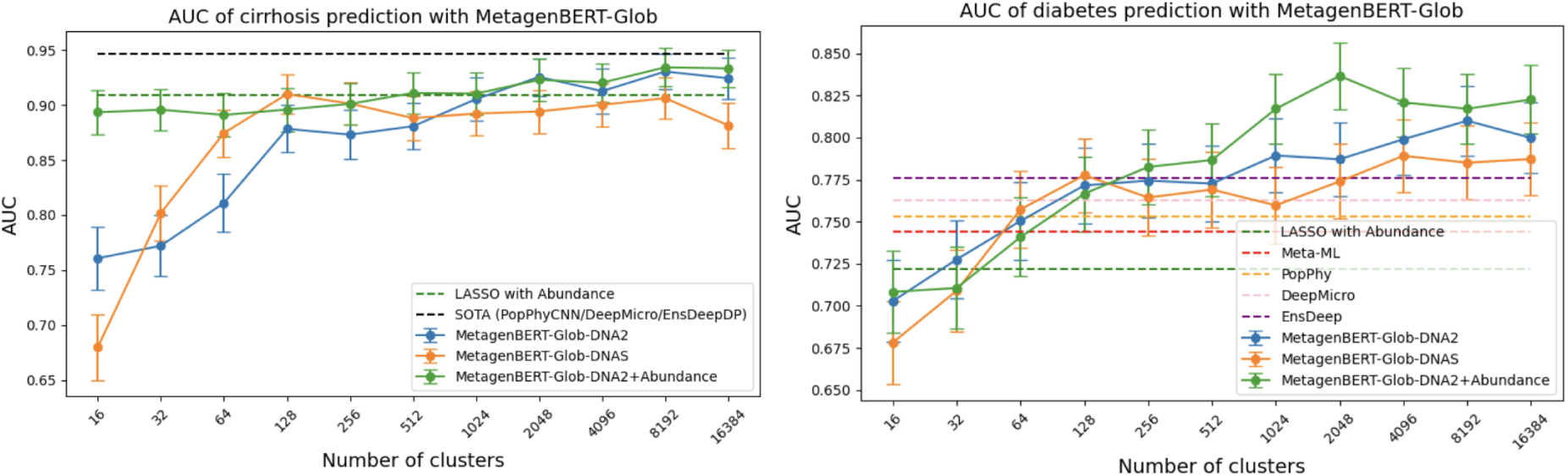
Performance Comparison between various SOTA models and Abundance, MetagenBERT-Glob and the concatenation of both with LASSO model. MetagenBERT-Glob results in better performance than Abundance alone and comparable to SOTA models when using a high enough number of clusters. Combining both abundance and global clustering results in better performance than each by themselves, supporting the idea that both carry different types of information. DNABERT-S embeddings result in better results than abundance alone, but fall short to DNABERT-2 performances, especially for the harder task of Diabetes prediction

### 3.3. Classification with a subsample of reads

We compare the results obtained using subsamples from our cirrhosis dataset to those obtained with the local clustering approach to assess the relevance of the clustering step. The subsampling method generally achieves poor results when using a small number of subsamples but improves as the number of reads increases. This is due to the fact that, when using a low number of samples„ there is a high risk of omitting important regions of the metagenome or overemphasizing non-relevant parts when the number of subsamples, this risk decreases when the number of subsamples increases.

A similar dynamic is observed in the local clustering results. However, the performance achieved with subsampling remains nonetheless lower, acknowledging the importance of the clustering step in our approach.

**Figure 8.**
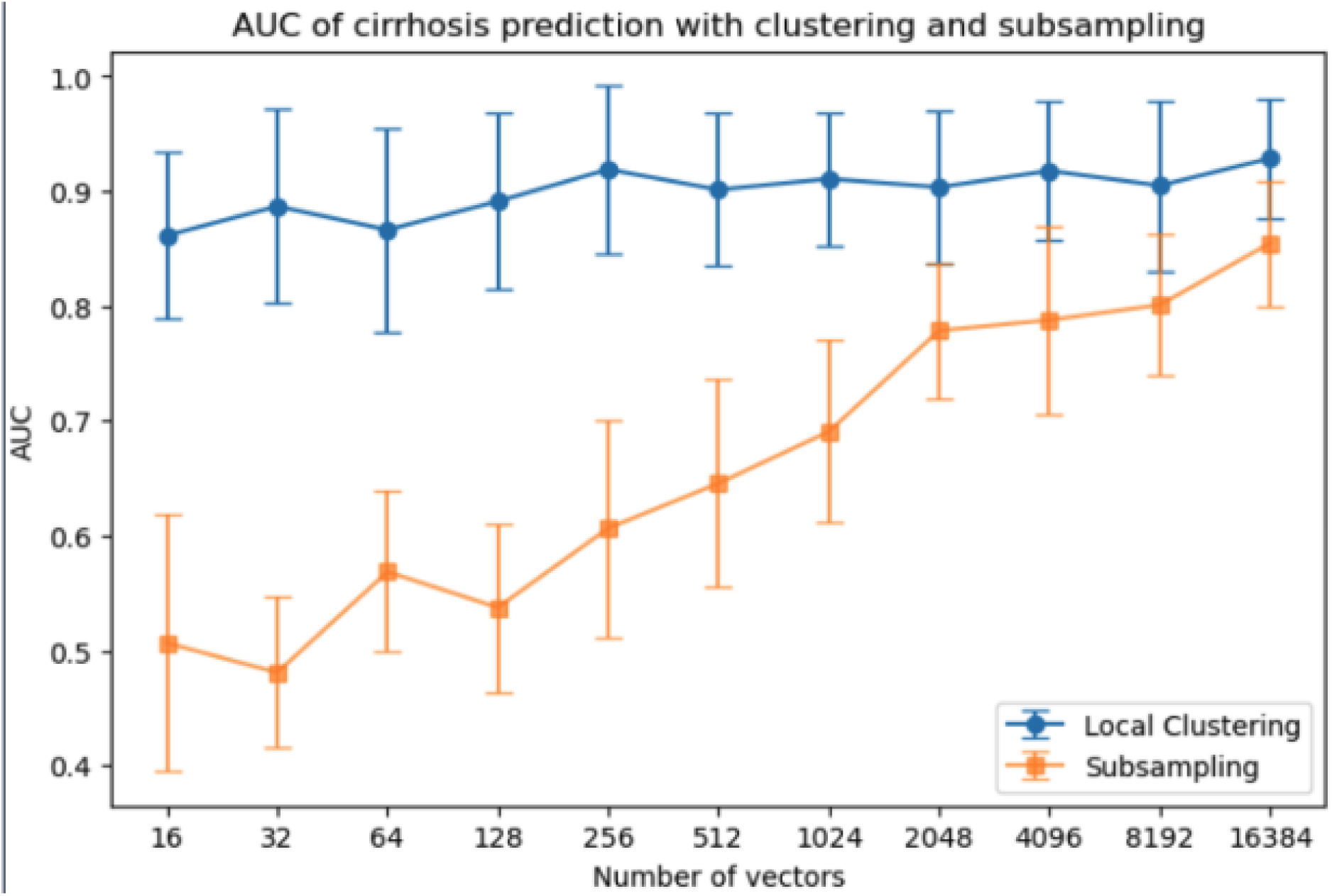
Performance Comparison between the local clustering method and the subsampling method. Although the performance increases with the size of the subset, we see that the local clustering method still outperforms this method, underlining the relevance of the clustering process

### 3.4. MetagenBERT-Glob remains accurate even when using a lower amount of data

As shown in Figure 9, MetagenBERT-Glob achieves robust performance even when applied to a fraction of the available read embeddings, without a substantial decrease in AUC. Across both datasets, and regardless of whether DNABERT-2 or DNABERT-S embeddings are used, results remain stable for every number of clusters. This resilience can be attributed to the compositional nature of the resulting representation: given the high number of reads per metagenomic sample, a representative subset is sufficient to preserve the integrity of the embedding. This property significantly enhances the scalability of our approach and reduces computational demands, making it more practical for large-scale metagenomic analyses.

**Figure 9.**
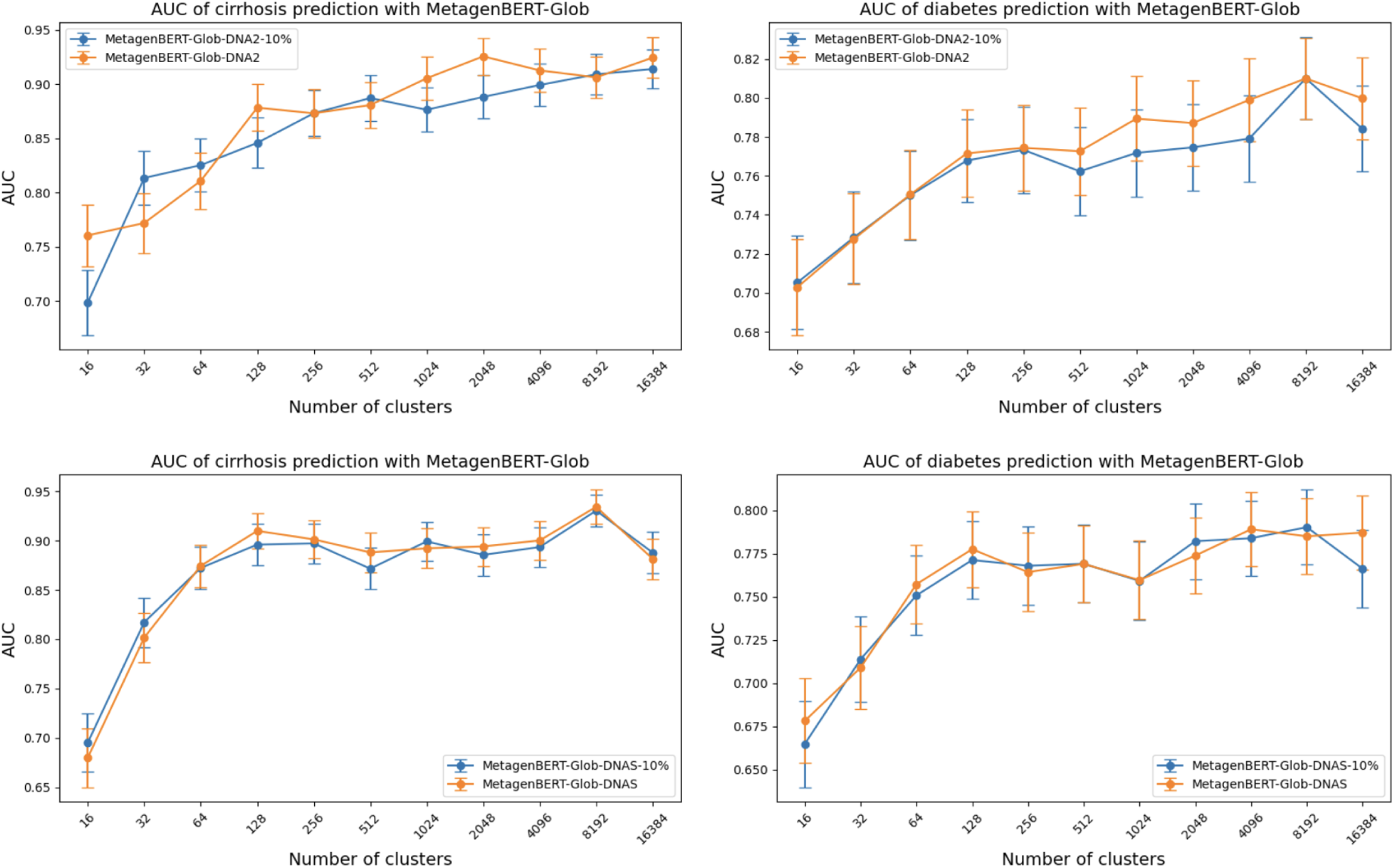
Comparison of Prediction performance when using MetagenBERT-Glob with all reads assigned and with only 10% assigned

### 3.5. Clusters analysis on a taxonomic and functional level

As we can see on Figure 10, analysis of the clusters identified by our MetagenBERT-Glob algorithm reveals variations in the mean entropy of individual clusters across different groups, exhibiting trends similar to those observed in the number of genomes identified using Centrifuge. Furthermore, a proportional relationship appears to exist between cluster entropy and the number of genomes retrieved within a sample. These observations support the notion that the clusters generated by our method capture meaningful microbiome dynamics, potentially reflecting underlying diversity and functional characteristics.

**Figure 10.**
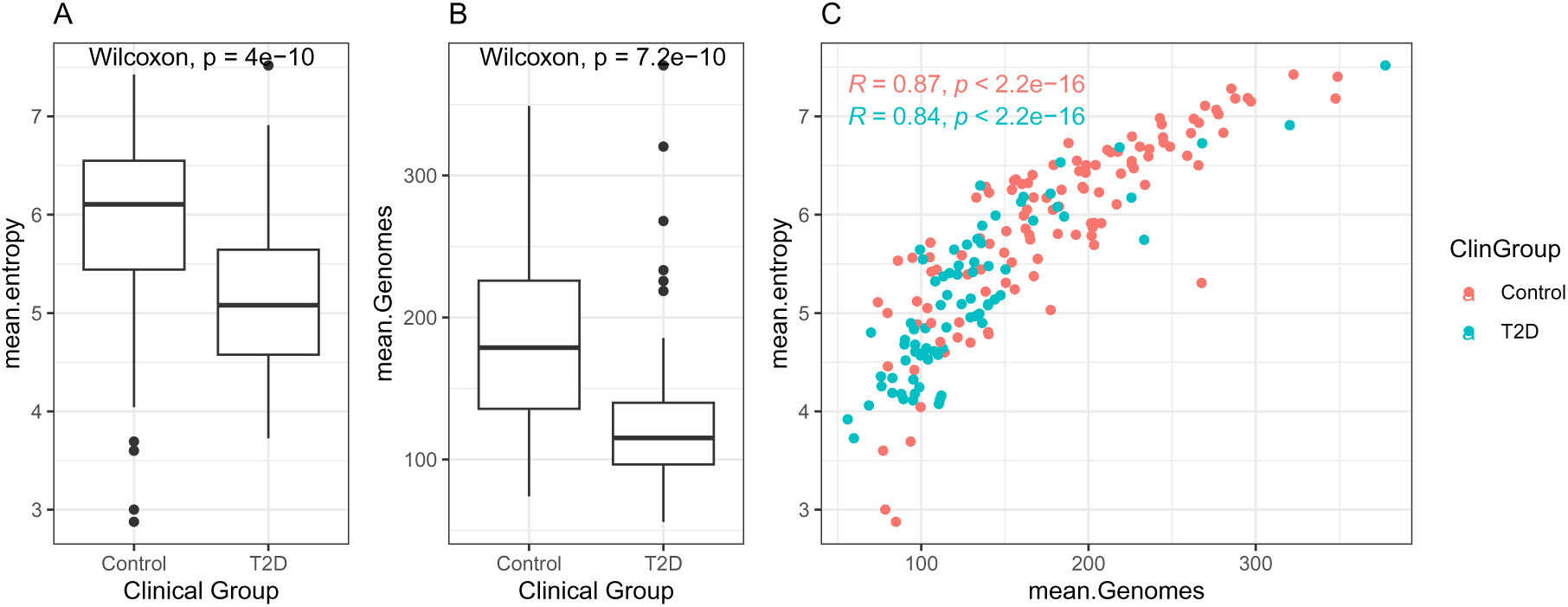
Analysis of Clusters composition obtained when using MetagenBERT-Glob with DNABERT-2 embeddings and 2048 clusters on Type 2 Diabetes (T2D) dataset. A) Mean entropy of clusters depending on their group. B) Mean number of genomes retrieved. C) Scatterplot of samples per group comparing their mean entropy and number of genomes retrieved

## 4. Discussion

In this study, we introduced MetagenBERT, an end-to-end, species-agnostic framework leveraging foundational DNA language models to classify diseases from raw metagenomic reads. Our results demonstrate that embedding millions of DNA reads with DNABERT-2 and DNABERT-S, followed by tailored aggregation strategies, yields classification performance comparable to or exceeding state-of-the-art models, especially in complex cases such as type 2 diabetes. Notably, the *local clustering* approach captures latent structure in the metagenome that appears more predictive than species abundance alone, suggesting that transformer-derived embeddings encode biologically meaningful features beyond taxonomy—potentially reflecting functional guilds, compositional biases, or strain-level genomic patterns.

Meanwhile, the MetagenBERT-Glob method constructs alternative abundance profiles that complement traditional species-based ones, with performance improving as the number of clusters increases. This synergy between taxonomic and embedding-based representations hints at orthogonal and biologically relevant information being captured by the transformer space. Importantly, the observed gains in prediction are not solely due to the expressive power of DNABERT-derived centroids, but also from the topology of the embedding space itself, as shown by the effectiveness of cluster-based abundance vectors.

While promising, our approach comes with limitations: the computational cost of embedding and clustering tens of millions of reads remains significant - although our results with only a fraction of the data show encouraging possibilities in simplifying the process by not embedding all the reads. Moreover, the interpretability of transformer embeddings is limited, and the datasets used—although standard—comprise relatively few samples, which could affect generalization. Future work will focus on scaling the method, improving biological interpretability (e.g. deeper functional annotation of clusters), and extending the framework to other microbiome types or multi-omics integration.

While Transformer-based models for read embedding, such as DNABERT-2, have demonstrated impressive performance, the field is rapidly evolving, with novel architectures and pretraining strategies emerging regularly that may yield even more informative embeddings. Notably, DNABERT-2 is trained on a diverse set of species, including humans, various animals, viruses, and fungi. This broad training scope, while valuable, may not optimally capture the unique characteristics of metagenomic sequences. Thus, there is potential in developing a model pre-trained specifically on metagenomic datasets, which could provide embeddings better suited for this domain.

We emphasize the complexity of our embedding space: each metagenome consists of tens of millions of reads, embedded here in a 768-dimensional vector space. For clustering, we initially employed the K-Means algorithm due to its simplicity and scalability. However, K-Means relies on Euclidean distance, which may encounter some issues in high-dimensional spaces due to the curse of dimensionality G., 1979. This phenomenon often causes distance metrics to lose their discriminative power, potentially impairing clustering performance. To address this limitation, we propose exploring more suitable alternatives for high-dimensional clustering. In particular, HDBSCAN Malzer and Baum, 2020, a hierarchical density-based algorithm, offers improved sensitivity to varying local densities and is generally more robust than K-Means in complex data landscapes. Additionally, subspace clustering techniques such as CLIQUE or PROCLUS “Automatic subspace clustering of high dimensional data for data mining applications” n.d. Aggarwal and Procopiuc, n.d. may uncover structure within meaningful low-dimensional subspaces of the embedding space. Spectral clustering Ng et al., n.d. also presents a compelling approach, especially when preceded by dimensionality reduction techniques such as PCA or UMAP. However, our tests in reducing dimension with PCA showed low variance when dividing dimension by a factor of 3 to 12, suggesting the dimension reduction loses many valuable features and information.

Furthermore, in the case of MetagenBERT-Local, we decided to address the Multiple Instance Learning problem with the DeepSets architecture. We want to point out that this architecture can face some issues like aggregation bottlenecks or lack of pairwise interaction. To better capture the structure of our clustered space, we suggest using more modern networks such as Set Transformers Lee et al., 2019, Graph Neural Networks Zhou et al., 2020 and PointNet++ Qi et al., 2017.

Additionally, we envision that MetagenBERT could serve as a building block for more interpretable and functionally grounded microbiome diagnostics, particularly in clinical contexts where species-level methods fall short. Overall, our results suggest that language-model-based embeddings represent a novel and promising axis of representation for metagenomic data, capable of enriching or even transcending conventional microbiome analysis pipelines.

## Data and Code Availability

DNA sequences are available from EBI (ERP005860) for cirrhosis and NCBI ( SRA045646 and SRA050230) for type 2 diabetes. The code can be found on Github following this link : https://github.com/CorvusVaine/MetagenBERT

## Acknowledgments

This work was granted access to the HPC resources of IDRIS under the allocations 2023-AD011014580, 2024-AD011014580R1 and 2024-AD011015723R1 made by GENCI

## References

Aggarwal CC, Procopiuc C (n.d.). Fast algorithms for projected clustering ().

AltschuP SF, Gish W, Miller W, Myers EW, Lipman DJ (n.d.). Basic Local Alignment Search Tool (), 8.

Automatic subspace clustering of high dimensional data for data mining applications (n.d.) ().

Babenko B (n.d.). Multiple Instance Learning: Algorithms and Applications ().

Calle ML (2019). Statistical Analysis of Metagenomics Data. Genomics & Informatics 17, e6. 10.5808/GI.2019.17.1.e6. URL: http://genominfo.org/journal/view.php?doi=10.5808/GI.2019.17.1.e6 (visited on 02/16/2023).

Chen S (2023). Ultrafast one-pass FASTQ data preprocessing, quality control, and deduplication using fastp. iMeta 2, e107. 10.1002/imt2 . 107. URL: https://onlinelibrary.wiley.com/doi/10.1002/imt2.107 (visited on 04/06/2025).

Dalla-Torre H, Gonzalez L, Mendoza-Revilla J, Lopez Carranza N, Grzywaczewski AH, Oteri F, Dallago C, Trop E, De Almeida BP, Sirelkhatim H, Richard G, Skwark M, Beguir K, Lopez M, Pierrot T (2025). Nucleotide Transformer: building and evaluating robust foundation models for human genomics. Nature Methods 22, 287–297. 10.1038/s41592-024-02523-z. URL: https://www.nature.com/articles/s41592-024-02523-z (visited on 04/06/2025).

Douze M, Guzhva A, Deng C, Johnson J, Szilvasy G, Mazaré PE, Lomeli M, Hosseini L, Jégou H (2024). The Faiss library. arXiv: 2401.08281 [cs.LG].

Ferranti EP, Dunbar SB, Dunlop AL, Corwin EJ (2014). 20 Things You Didn’t Know About the Human Gut Microbiome. Journal of Cardiovascular Nursing 29, 479–481. 10.1097/JCN.0000000000000166. URL: https://journals.lww.com/00005082-201411000-00004 (visited on 04/06/2025).

G. T (1979). A Problem of Dimensionality: A Simple Example. IEEE Transactions on Pattern Analysis and Machine Intelligence. 10.1109/TPAMI.1979.4766926.

Johnson J, Douze M, Jégou H (2019). Billion-scale similarity search with GPUs. IEEE Transactions on Big Data 7, 535–547.

Le Chatelier E, Nielsen T, Qin J, Prifti E, Hildebrand F, Falony G, Almeida M, Arumugam M, Batto JM, Kennedy S, Leonard P, Li J, Burgdorf K, Grarup N, Jørgensen T, Brandslund I, Nielsen HB, Juncker AS, Bertalan M, Levenez F, et al. (2013). Richness of human gut microbiome correlates with metabolic markers. Nature 500, 541–546. 10.1038/nature12506. URL: https://www.nature.com/articles/nature12506 (visited on 12/05/2022).

Lee J, Lee Y, Kim J, Kosiorek AR, Choi S, Teh YW (2019). Set Transformer: A Framework for Attention-based Permutation-Invariant Neural Networks. arXiv:1810.00825 [cs]. 10.48550/arXiv.1810.00825. URL: http://arxiv.org/abs/1810.00825 (visited on 04/06/2025).

Lee SJ, Rho M (2022). Multimodal deep learning applied to classify healthy and disease states of human microbiome. Scientific Reports 12, 824. 10.1038/s41598-022-04773-3. URL: https://www.nature.com/articles/s41598-022-04773-3 (visited on 07/25/2022).

Liang Q, Bible PW, Liu Y, Zou B, Wei L (2020). DeepMicrobes: taxonomic classification for metagenomics with deep learning. NAR Genomics and Bioinformatics 2, lqaa009. 10.1093/nargab/lqaa009. URL: https://academic.oup.com/nargab/article/doi/10.1093/nargab/lqaa009/5740226 (visited on 06/22/2022).

Malzer C, Baum M (2020). A Hybrid Approach To Hierarchical Density-based Cluster Selection. In: 2020 *IEEE International Conference on Multisensor Fusion and Integration for Intelligent Systems (MFI)*. arXiv:1911.02282 [cs], pp. 223–228. 10.1109/MFI49285.2020.9235263. URL: http://arxiv.org/abs/1911.02282 (visited on 04/06/2025).

MetaHIT Consortium, Qin J, Li R, Raes J, Arumugam M, Burgdorf KS, Manichanh C, Nielsen T, Pons N, Levenez F, Yamada T, Mende DR, Li J, Xu J, Li S, Li D, Cao J, Wang B, Liang H, Zheng H, et al. (2010). A human gut microbial gene catalogue established by metagenomic sequencing. Nature 464, 59–65. 10.1038/nature08821. URL: http://www.nature.com/articles/nature08821 (visited on 02/14/2023).

Mock F, Kretschmer F, Kriese A, Böcker S, Marz M (2021). BERTax: taxonomic classification of DNA sequences with Deep Neural Networks. preprint. Bioinformatics. 10.1101/2021.07.09.451778. URL: http://biorxiv.org/lookup/doi/10.1101/2021.07.09.451778 (visited on 05/11/2022).

Mulenga M, Abdul Kareem S, Qalid Md Sabri A, Seera M, Govind S, Samudi C, Mohamad SB (2021). Feature Extension of Gut Microbiome Data for Deep Neural Network-Based Colorectal Cancer Classification. IEEE Access 9, 23565–23578. 10.1109/ACCESS.2021.3050838. URL: https://ieeexplore.ieee.org/document/9319639/ (visited on 07/28/2022).

Ng AY, Jordan MI, Weiss Y (n.d.). On Spectral Clustering: Analysis and an algorithm ().

Nissen JN, Johansen J, Allesøe RL, Sønderby CK, Armenteros JJA, Grønbech CH, Jensen LJ, Nielsen HB, Petersen TN, Winther O, Rasmussen S (2021). Improved metagenome binning and assembly using deep variational autoencoders. Nature Biotechnology 39, 555–560. 10.1038/s41587-020-00777-4. URL: http://www.nature.com/articles/s41587-020-00777-4 (visited on 08/30/2022).

Oehler JB, Wright H, Stark Z, Mallett AJ, Schmitz U (2023). The application of long-read sequencing in clinical settings. Human Genomics 17, 73. 10.1186/s40246-023-00522-3. URL: https://humgenomics.biomedcentral.com/articles/10.1186/s40246-023-00522-3 (visited on 04/06/2025).

Oh M, Zhang L (2020). DeepMicro: deep representation learning for disease prediction based on microbiome data. Scientific Reports 10, 6026. 10.1038/s41598-020-63159-5. URL: http://www.nature.com/articles/s41598-020631595 (visited on 07/26/2022).

Pasolli E, Truong DT, Malik F, Waldron L, Segata N (2016). Machine Learning Meta-analysis of Large Metagenomic Datasets: Tools and Biological Insights. PLOS Computational Biology 12. Ed. by Jonathan A. Eisen, e1004977. 10.1371/journal.pcbi.1004977. URL: https://dx.plos.org/10.1371/journal.pcbi.1004977 (visited on 09/12/2022).

Pflughoeft KJ, Versalovic J (2012). Human Microbiome in Health and Disease. Annual Review of Pathology: Mechanisms of Disease 7, 99–122. 10.1146/annurev-pathol-011811-132421. URL: https://www.annualreviews.org/doi/10.1146/annurev-pathol-011811-132421 (visited on 10/16/2022).

Portes J, Trott A, Havens S, King D, Venigalla A, Nadeem M, Sardana N, Khudia D, Frankle J (n.d.). MosaicBERT: A Bidirectional Encoder Optimized for Fast Pretraining ().

Qi CR, Yi L, Su H, Guibas LJ (2017). PointNet++: Deep Hierarchical Feature Learning on Point Sets in a Metric Space. arXiv:1706.02413 [cs]. 10.48550/arXiv.1706.02413. URL: http://arxiv.org/abs/1706.02413 (visited on 04/06/2025).

Qin N, Yang F, Li A, Prifti E, Chen Y, Shao L, Guo J, Le Chatelier E, Yao J, Wu L, Zhou J, Ni S, Liu L, Pons N, Batto JM, Kennedy SP, Leonard P, Yuan C, Ding W, Chen Y, et al. (2014). Alterations of the human gut microbiome in liver cirrhosis. Nature 513, 59–64. 10.1038/nature13568. URL: http://www.nature.com/articles/nature13568 (visited on 02/22/2023).

Queyrel M, Prifti E, Templier A, Zucker JD (2020). Towards end-to-end disease prediction from raw metagenomic data. preprint. Genomics. 10.1101/2020.10.29.360297. URL: http://biorxiv.org/lookup/doi/10.1101/2020.10.29.360297 (visited on 09/09/2022).

Reiman D, Metwally AA, Sun J, Dai Y (2020). PopPhy-CNN: A Phylogenetic Tree Embedded Architecture for Convolutional Neural Networks to Predict Host Phenotype From Metagenomic Data. IEEE Journal of Biomedical and Health Informatics 24, 2993–3001. 10.1109/JBHI.2020.2993761. URL: https://ieeexplore.ieee.org/document/9091025/ (visited on 07/25/2022).

Roy G, Prifti E, Belda E, Zucker JD (2024). Deep learning methods in metagenomics: a review. Microbial Genomics 10. 10.1099/mgen.0.001231. URL: https://www.microbiologyresearch.org/content/journal/mgen/10.1099/mgen.0.001231 (visited on 04/06/2025).

Sczyrba A, Hofmann P, Belmann P, Koslicki D, Janssen S, Dröge J, Gregor I, Majda S, Fiedler J, Dahms E, Bremges A, Fritz A, Garrido-Oter R, Jørgensen TS, Shapiro N, Blood PD, Gurevich A, Bai Y, Turaev D, DeMaere MZ, et al. (2017). Critical Assessment of Metagenome Interpretation—a benchmark of metagenomics software. Nature Methods 14, 1063–1071. 10.1038/nmeth.4458. URL: http://www.nature.com/articles/nmeth.4458 (visited on 10/17/2022).

Sharma D, Paterson AD, Xu W (2020). TaxoNN: ensemble of neural networks on stratified microbiome data for disease prediction. Bioinformatics 36. Ed. by Pier Luigi Martelli, 4544–4550. 10.1093/bioinformatics/btaa542. URL: https://academic.oup.com/bioinformatics/article/36/17/4544/5843784 (visited on 09/09/2022).

Shen Y, Zhu J, Deng Z, Lu W, Wang H (2022). Ensdeepdp: An Ensemble Deep Learning Approach for Disease Prediction Through Metagenomics. IEEE/ACM Transactions on Computational Biology and Bioinformatics, 1–14. 10.1109/TCBB.2022.3201295. URL: https://ieeexplore.ieee.org/document/9866523/ (visited on 08/31/2022).

Thomas AM, Segata N (2019). Multiple levels of the unknown in microbiome research. BMC Biology 17, 48. 10.1186/s12915-019-0667-z. URL: https://bmcbiol.biomedcentral.com/articles/10.1186/s12915-019-0667-z (visited on 04/06/2025).

Wichmann A, Buschong E, Müller A, Jünger D, Hildebrandt A, Hankeln T, Schmidt B (2023). Meta-Transformer: deep metagenomic sequencing read classification using self-attention models. NAR Genomics and Bioinformatics 5, lqad082. 10.1093/nargab/lqad082. URL: https://academic.oup.com/nargab/article/doi/10.1093/nargab/lqad082/7269178 (visited on 04/06/2025).

Wu G, Zhao N, Zhang C, Lam YY, Zhao L (2021). Guild-based analysis for understanding gut microbiome in human health and diseases. Genome Medicine 13, 22. 10.1186/s13073-021-00840-y. URL: https://genomemedicine.biomedcentral.com/articles/10.1186/s13073-021-00840-y (visited on 06/22/2022).

Zaheer M, Kottur S, Ravanbakhsh S, Poczos B, Salakhutdinov R, Smola A (2018). Deep Sets. arXiv:1703.06114 [cs]. 10.48550/arXiv.1703.06114. URL: http://arxiv.org/abs/1703.06114 (visited on 04/06/2025).

Zhou J, Cui G, Hu S, Zhang Z, Yang C, Liu Z, Wang L, Li C, Sun M (2020). Graph neural networks: A review of methods and applications. AI Open 1, 57–81. 10.1016/j.aiopen.2021.01.001. URL: https://linkinghub.elsevier.com/retrieve/pii/S2666651021000012 (visited on 04/06/2025).

Zhou Z, Ji Y, Li W, Dutta P, Davuluri R, Liu H (2024a). DNABERT-2: Efficient Foundation Model and Benchmark For Multi-Species Genome. arXiv:2306.15006 [q-bio]. 10.48550/arXiv.2306.15006. URL: http://arxiv.org/abs/2306.15006 (visited on 04/06/2025).

Zhou Z, Wu W, Ho H, Wang J, Shi L, Davuluri RV, Wang Z, Liu H (2024b). DNABERT-S: Pioneering Species Differentiation with Species-Aware DNA Embeddings. arXiv:2402.08777 [q-bio]. 10.48550/arXiv.2402.08777. URL: http://arxiv.org/abs/2402.08777 (visited on 04/06/2025).

